## Supplementary Figures 1-5 for "The actomyosin system is essential for the integrity of the endosomal system in bloodstream form *Trypanosoma brucei*"

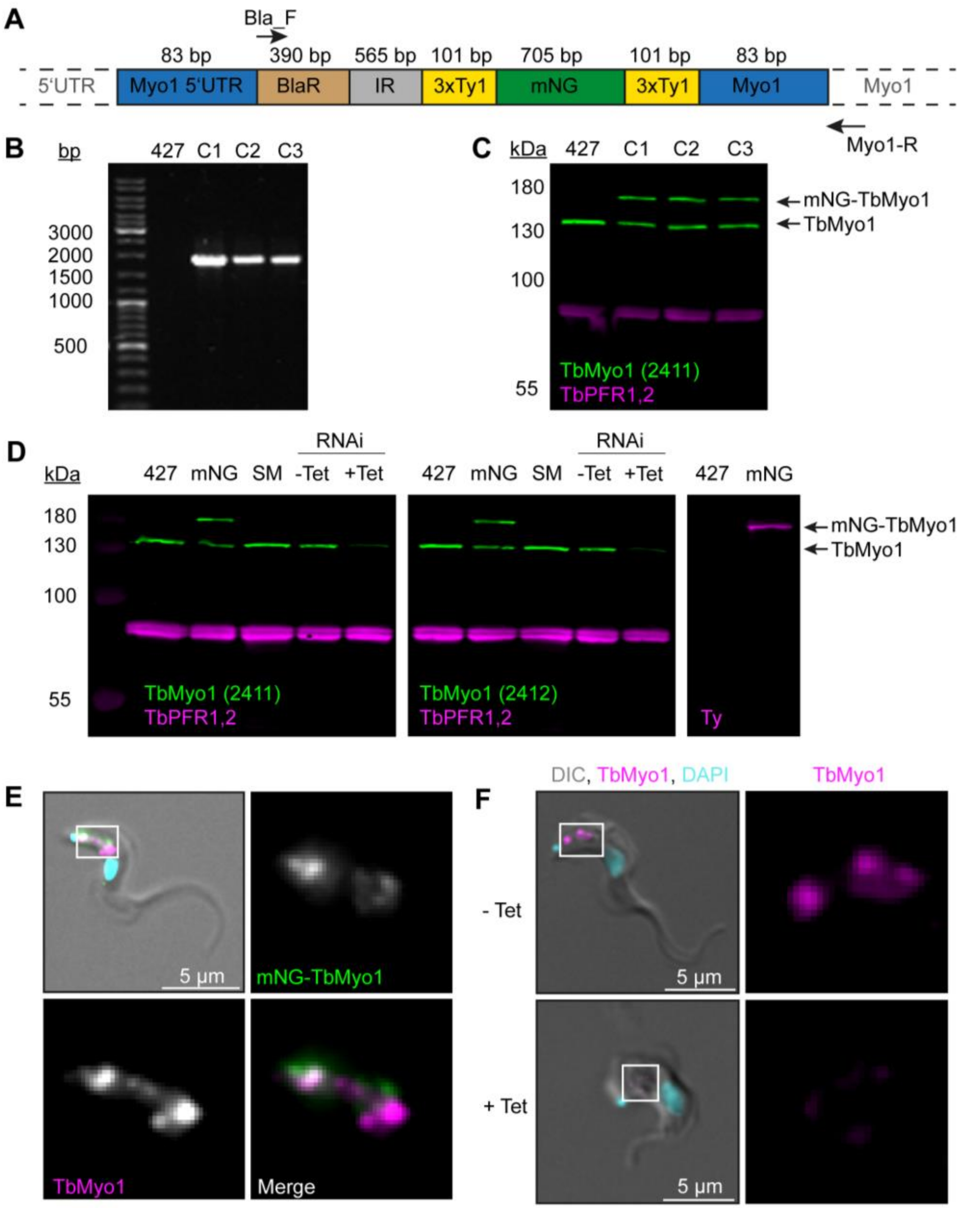
A
Bla_F
83 bp
390 bp
565 bp
101 bp
705 bp
101 bp
83 bp
5'UTR
Myo1 5'UTR
BlaR
IR
3xTy1
mNG
3xTy1
Myo1
Myo1
Myo1-R
B
bp
427 C1 C2 C3
3000
2000
1500
1000
500
C
kDa
427 C1 C2 C3
180
130
100
55
mNG-TbMyo1
TbMyo1
TbMyo1 (2411)
TbPFR1,2
D
kDa
RNAi
427 mNG SM -Tet +Tet
RNAi
427 mNG SM -Tet +Tet
427 mNG
180
130
100
55
mNG-TbMyo1
TbMyo1
TbMyo1 (2411)
TbPFR1,2
TbMyo1 (2412)
TbPFR1,2
Ty
E
5 µm
mNG-TbMyo1
TbMyo1
Merge
F
DIC, TbMyo1, DAPI
TbMyo1
- Tet
5 µm
+ Tet
5 µm

**Figure S1. Generation and validation of TbMyo1 antibodies and cell lines. (A)** Schematic of the TbMyo1 *in situ* tagging construct integrated into the genome. The construct encoded the 3' end of the Tb*MYO1* 5' UTR, a blasticidin (BlaR) resistance gene, the alpha/beta tubulin intergenic region (IR), the mNG gene flanked by two 3xTy1 epitope tags, and the 5' end of the Tb*MYO1* ORF. The sizes of the various elements are shown in the schematic. The 83 bp labels indicate the homology arms that drive homologous recombination at the *MYO1* locus. The annealing sites of the two primers used to confirm integration at the endogenous *MYO1* locus (Bla-F, Myo1-R) are indicated with arrows. Note that the schematic elements are not shown to scale. **(B)** Correct integration of the TbMyo1 tagging construct at the endogenous locus. The gDNA of wild-type (427) and 3 candidate mNG-TbMyo1 clones (C1 - C3) was purified and analysed by PCR. The annealing sites of the PCR primers are shown in panel A above. No product was observed for the wild-type control. An ~ 1700 bp product was obtained from the three mNG-TbMyo1 candidates, indicating correct integration. **(C)** Expression of mNG-TbMyo1. Whole-cell lysates from the three mNG-TbMyo1 clones and a wild-type control were prepared and analysed by immunoblotting. Anti-TbMyo1(2411) was used as a primary antibody; anti-TbPFR1,2 (L13D6) antibodies were used to detect the housekeeping proteins PFR1,2. The endogenous TbMyo1 signal (green) was visible in all lanes at ~130 kDa; C1, C2 and C3 additionally expressed an ~ 170 kDa protein, matching the size of the mNG-TbMyo1. **(D)** Validation of anti-TbMyo1 specificity. Whole cell lysates from four different cell lines (wild-type (427), single marker (SM), mNG-TbMyo1 (mNG), TbMyo1 RNAi) were immunoblotted with two anti-TbMyo1 antisera (2411, 2412). The TbMyo1 RNAi lysates were obtained from uninduced (-Tet) and tetracycline-induced (+Tet) samples. TbPFR1,2 was used as a loading control. Both antisera were able to detect the endogenous TbMyo1 protein at ~ 130 kDa. The signal intensities of the 427 cells and the single marker (SM) ones were almost equal, independent of which antiserum was used. An additional band at ~ 170 kDa was observed in the mNG-TbMyo1 samples. This upper band could also be detected using anti-Ty1 antibodies (right-hand image). In the RNAi-induced (+ Tet) cells, the signal from both antisera was weaker. **(E)** Strong overlap between endogenous and mNG-tagged TbMyo1. The mNG-TbMyo1 clones were fixed, labelled with anti-TbMyo1 antibodies (magenta), and imaged using widefield microscopy. DNA (cyan) was stained using DAPI. Both the mNG-TbMyo1 (green) and the endogenous protein were exclusively localised to the posterior region of the cells. Identical results were obtained using anti-Ty1 antibodies to label the mNG-TbMyo1. The same pattern was observed in all three clones; an exemplary image is shown. **(F)** Depletion of TbMyo1 results in a loss of signal and morphological changes. TbMyo1 RNAi cells were induced for 48h, fixed, and labelled with anti-TbMyo1 antibodies (magenta). DNA was stained with DAPI (cyan). Control cells showed an intense signal of TbMyo1 (magenta) between the nucleus and the kinetoplast. Induced cells showed a nearly complete loss of the signal, with a small degree of clone-to-clone variation. The morphology of the TbMyo1-depleted cells changed dramatically, with some cells developing a ‘Big Eye’ phenotype. The same results were obtained with three separate clones; an exemplary image is shown.

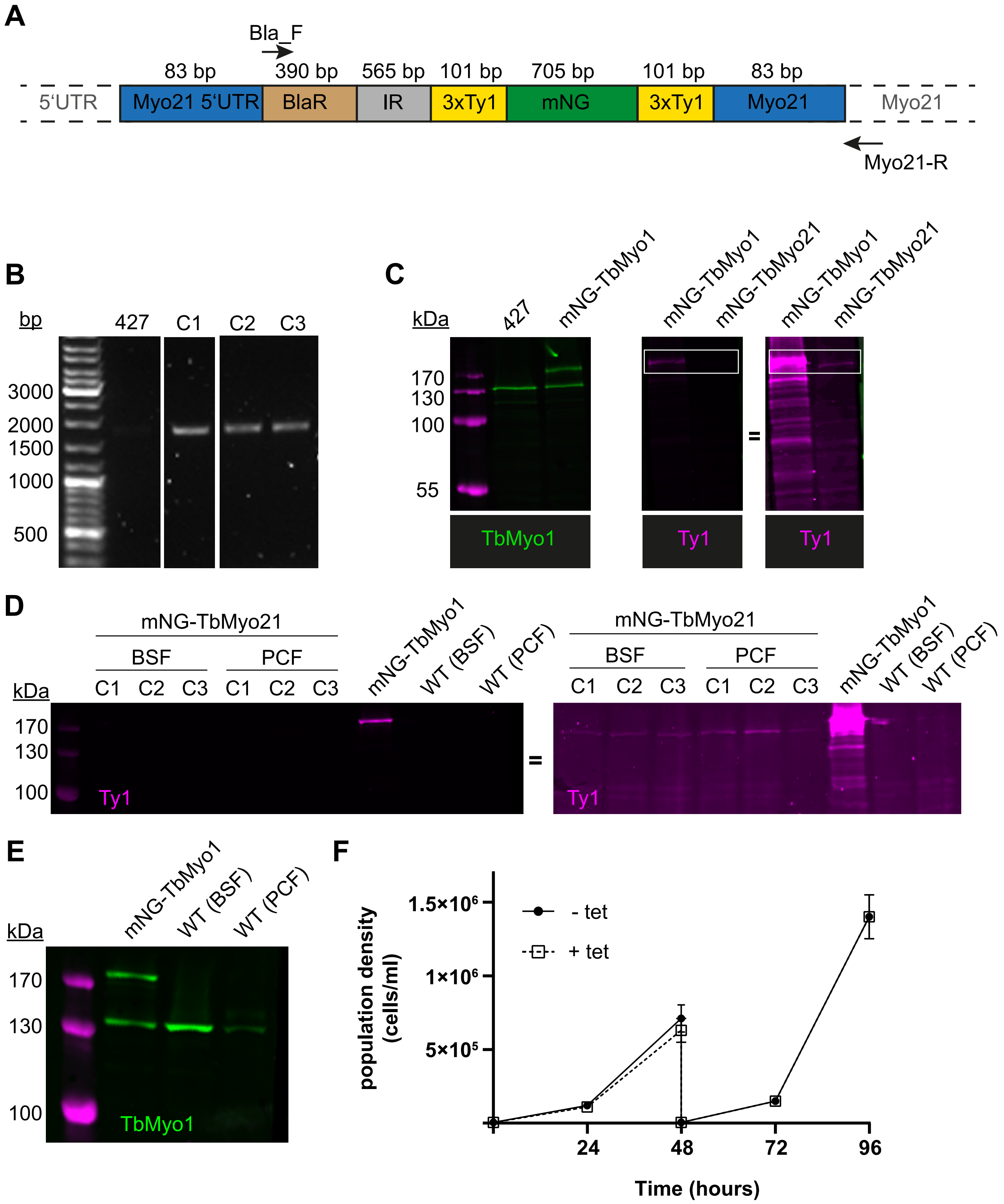
A
Bla_F
83 bp
390 bp
565 bp
101 bp
705 bp
101 bp
83 bp
5'UTR
Myo21 5'UTR
BlaR
IR
3xTy1
mNG
3xTy1
Myo21
Myo21
Myo21-R
B
bp
427
C1
C2
C3
3000
2000
1500
1000
500
C
kDa
427
mNG-TbMyo1
mNG-TbMyo1
mNG-TbMyo21
mNG-TbMyo1
mNG-TbMyo21
170
130
100
55
=
TbMyo1
Ty1
Ty1
D
mNG-TbMyo21
BSF
PCF
C1
C2
C3
C1
C2
C3
mNG-TbMyo1
WT (BSF)
WT (PCF)
kDa
170
130
100
Ty1
=
mNG-TbMyo21
BSF
PCF
C1
C2
C3
C1
C2
C3
mNG-TbMyo1
WT (BSF)
WT (PCF)
Ty1
E
kDa
mNG-TbMyo1
WT (BSF)
WT (PCF)
170
130
100
TbMyo1
F
population density (cells/ml)
1.5×10⁶
1×10⁶
5×10⁵
- tet
+ tet
24
48
72
96
Time (hours)

**Figure S2. TbMyo21 is expressed at an extremely low level. (A)** Generation of mNG-tagged TbMyo21 cell line. Schematic of the TbMyo21 *in situ* tagging construct integrated into the genome. The construct encoded the 3' end of the Tb*MYO21* 5' UTR, a blasticidin (BlaR) resistance gene, the alpha/beta tubulin intergenic region (IR), the mNG gene flanked by two 3xTy1 epitope tags, and the 5' end of the Tb*MYO21* ORF. The sizes of the various elements are shown in the schematic. The 83 bp labels indicate the homology arms that drive homologous recombination at the *MYO21* locus. The annealing sites of the primers used to confirm integration at the endogenous *MYO21* locus (Bla-F, Myo21-R) are indicated with arrows. Note that the schematic elements are not shown to scale. **(B)** Correct integration of the TbMyo21 tagging construct at the endogenous locus. The gDNA of wild-type (427) and 3 candidate mNG-TbMyo1 clones (C1 - C3) was purified and analysed by PCR. The annealing sites of the PCR primers are shown in panel A. No product was observed for the wild-type control (427). An ~ 2000 bp product was obtained from the three mNG-TbMyo21 candidates, indicating correct integration. **(C)** mNG-TbMyo21 is expressed at a very low level relative to TbMyo1. Whole-cell lysates from a wild-type control, the mNG-TbMyo1 cell line, and the mNG-TbMyo21 cell line were prepared and analysed by immunoblotting. The endogenous TbMyo1 signal (green) was visible at ~130 kDa; the mNG-TbMyo1 cell line additionally expressed an ~ 170 kDa protein, corresponding to mNG-TbMyo1. The anti-Ty1 antibody signal (magenta) required an overexposure (right hand image) to visualise the mNG-TbMyo21 band at ~ 160 kDa (boxed area). **(D)** TbMyo21 expression levels are equally low in bloodstream form (BSF) and procyclic form (PCF) cells. Whole-cell lysates from the mNG-TbMyo21 cell line and a wild-type control were prepared before and after differentiation to PCF cells and analysed by immunoblotting. A whole-cell lysate from BSF mNG-TbMyo1 cells served as a positive control. The anti-Ty1 antibody signal (magenta) was only visible in the mNG-TbMyo1 cell line at ~ 170 kDa (left-hand image). The anti-Ty1 signal required an overexposure to visualise an mNG-TbMyo21 band at ~ 160 kDa (right-hand image). **(E)** TbMyo1 expression is downregulated after differentiation to PCF cells. The whole-cell lysates from BSF mNG-TbMyo1 cells and both BSF and PCF wild-type cells were analysed by immunoblotting. Equal numbers of cell equivalents ($1.4 \times 10^6$) were loaded in each lane. The endogenous TbMyo1 signal (green) was visible in all lanes at ~130 kDa; the mNG-TbMyo1 cell line additionally expressed an ~ 170 kDa protein, matching the size of the mNG-TbMyo1. Quantification was by direct measurement of band intensity after background subtraction and without normalisation to a loading control, and is therefore somewhat tentative. The TbMyo1 signal in PCF cells showed <50% signal intensity compared to BSF cells. **(F)** TbMyo21 does not appear to be essential for growth *in vitro*. The cell population density of TbMyo21 RNAi cells was measured for 96 hours in tetracycline uninduced (-Tet) and induced (+Tet) conditions. The cells were diluted after 48 hours to ensure logarithmical growth. Results from three independent experiments using three separate clones are shown; data points are mean +/- standard deviation.

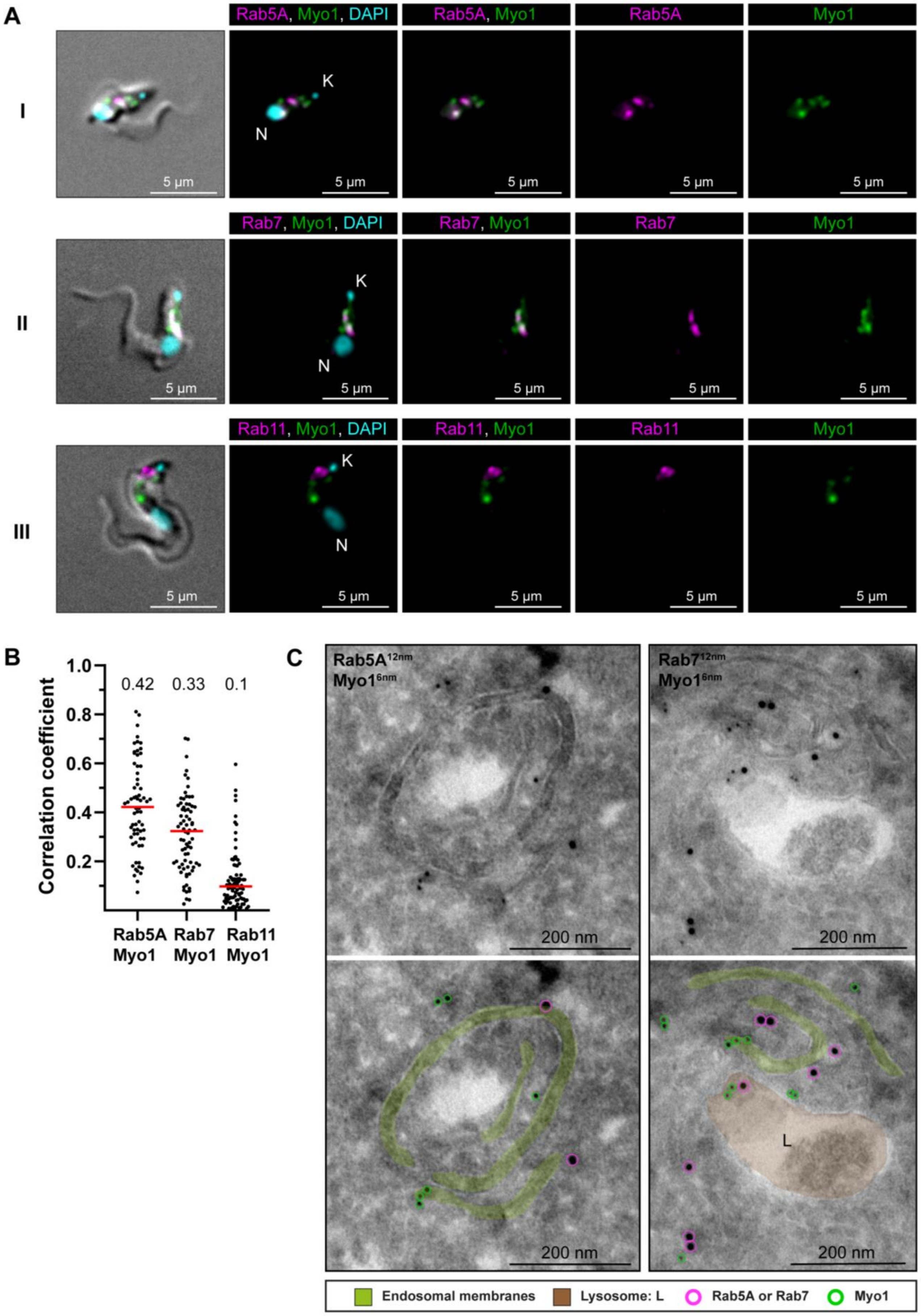
A
I
Rab5A, Myo1, DAPI
Rab5A, Myo1
Rab5A
Myo1
K
N
5 µm
II
Rab7, Myo1, DAPI
Rab7, Myo1
Rab7
Myo1
III
Rab11, Myo1, DAPI
Rab11, Myo1
Rab11
Myo1
B
Correlation coefficient
1.0
0.8
0.6
0.4
0.2
0.42
0.33
0.1
Rab5A Myo1
Rab7 Myo1
Rab11 Myo1
C
Rab5A12nm Myo16nm
Rab712nm Myo16nm
200 nm
L
Endosomal membranes
Lysosome: L
Rab5A or Rab7
Myo1

**Figure S3. TbMyo1 is associated with early and late endosomes. (A)** TbMyo1 signals partially overlap with TbRab5A and TbRab7 signals, but not with TbRab11 signals. Colocalisation experiments were conducted using chemically fixed mNG-TbMyo1 cells, labelled with antibodies against TbRab5A, 7, or 11 and imaged using widefield microscopy. The fluorescence signals for TbRab5A (I, magenta), TbRab7 (II, magenta), TbRab11 (III, magenta) and TbMyo1 (I – III, green) were localised between the kinetoplast (K) and the nucleus (N) of the cells. DNA was stained using DAPI (I – III, cyan). Exemplary cells from four independent experiments are shown. **(B)** Quantification of correlation between TbMyo1 and TbRab signals. Correlation was estimated using Pearson's correlation coefficient (TbRab5A/TbMyo1, n = 66; TbRab7/TbMyo1, n = 73; TbRab11/TbMyo1, n = 86). Red bars indicate the median R; median R values for each pair are shown above the dot plot. **(C)** Electron micrographs of cryosections colabelled with anti-TbMyo1 and anti-TbRab5A or anti-TbRab7 antibodies. For each panel a raw and a pseudocoloured version of the image are presented. Endosomal membranes are coloured green. The lysosome is coloured brown. Gold particles corresponding to TbMyo1 signals are outlined in green. Gold particles corresponding to TbRab5A or TbRab7 signals are outlined in magenta. Exemplary cells from a single labelling experiment are shown.

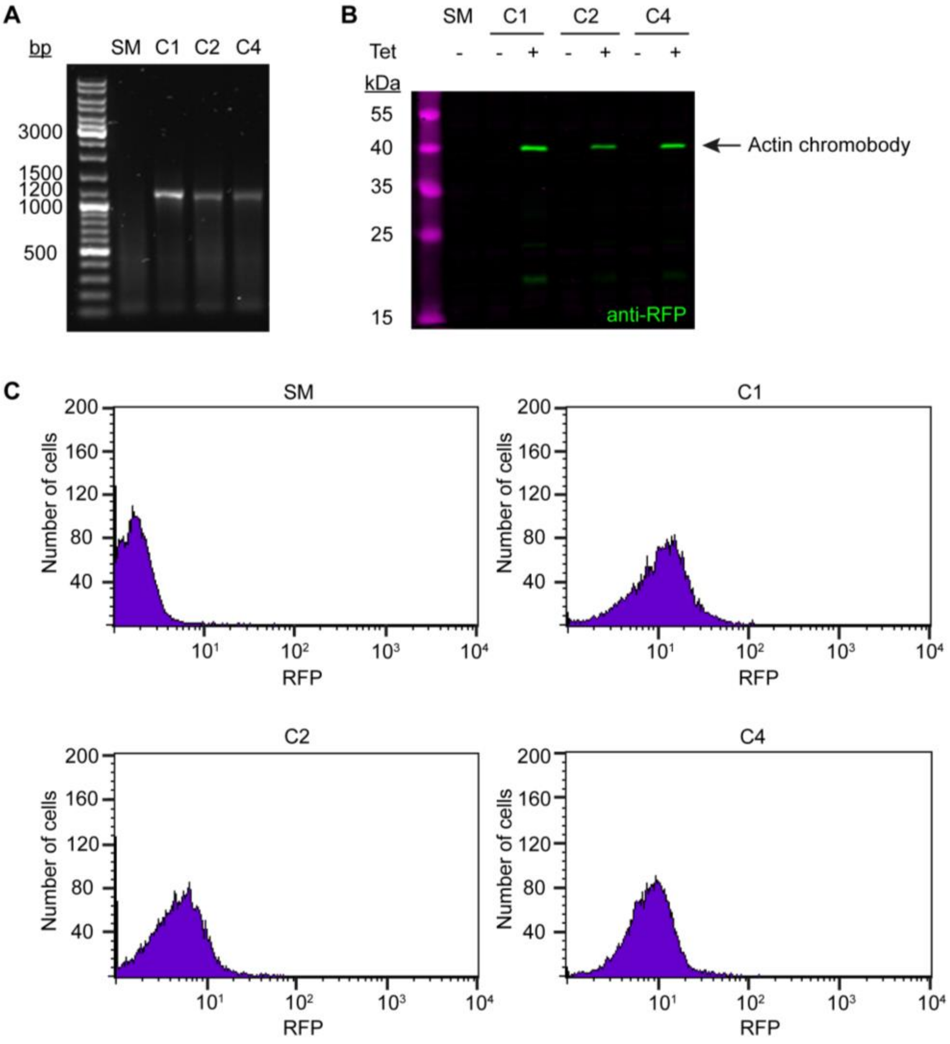


**Figure S4. Generation and validation of the anti-actin chromobody cell line. (A)** Integration of the anti-actin TagRFP chromobody construct. The gDNA of wild-type (SM) and 3 candidate clones (C1, C2, C4) was purified and analysed by PCR using primers specific for the chromobody. No product was observed for the wild-type control. An ~ 1200 bp product was obtained from the three candidates, indicating integration into the genome. **(B)** Tight and inducible expression of the anti-actin TagRFP chromobody. Whole-cell lysates from the three clones (tetracycline-induced and -uninduced) and a wild-type control (SM) were prepared and analysed by immunoblotting, using anti-RFP antibodies. No signal was detected in the control sample and in the uninduced (-) chromobody clones. In the induced (+) samples, a protein band at ~40 kDa was detected in all three clones. The expected size of the anti-actin chromobody is 41.5 kDa. **(C)** Varying anti-actin TagRFP chromobody expression levels in the three clones. The three clones (C1, C2, C4) were analysed 24 hours after induction with tetracycline and RFP signal was measured using flow cytometry. Wild-type cells (SM) served as a control. All clones showed a significant increase in fluorescence counts in the RFP channel relative to control cells, with some clone-to-clone variation in expression levels.

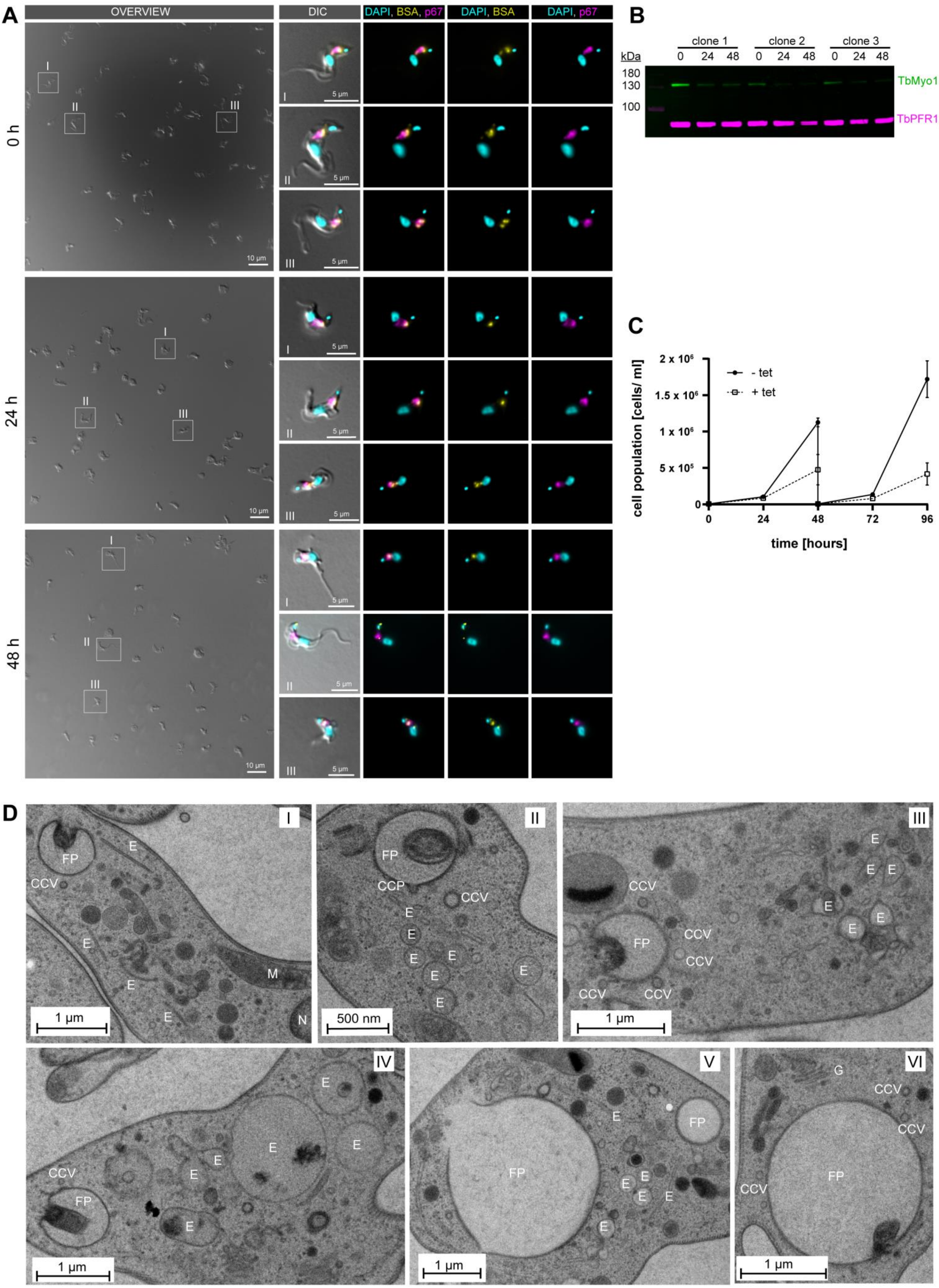
A
OVERVIEW
DIC
DAPI, BSA, p67
DAPI, BSA
DAPI, p67
0 h
24 h
48 h
I
II
III
10 µm
5 µm
B
clone 1
clone 2
clone 3
kDa
0 24 48
180
130
100
TbMyo1
TbPFR1,2
C
cell population [cells/ ml]
time [hours]
- tet
+ tet
2 x 10^6
1.5 x 10^6
1 x 10^6
5 x 10^5
0
0 24 48 72 96
D
IV
V
VI
FP
CCV
CCP
E
M
N
G
1 µm
500 nm

**Figure S5. TbMyo1 depletion affects cell morphology and endosomal membrane organisation. (A)** Bovine serum albumin (BSA) is still internalised after TbMyo1 depletion. TbMyo1 depletion was induced by RNAi for 0, 24, and 48 hours. The cells were then harvested and incubated with fluorophore-conjugated BSA (yellow) for 30 minutes at 37°C. The cells were then fixed with formaldehyde and immunolabeled with anti-p67 antibodies (magenta), and DNA was stained with DAPI (cyan). Exemplary DIC fields of view are presented alongside enlarged views of selected morphologically normal cells (I-III). Prolonged protein depletion led to an increased number of cells displaying a rounded morphology. In cells with normal morphology, fluorescent BSA strongly overlapped with p67, indicating normal cargo uptake. Results obtained in a single experiment using three separate TbMyo1 RNAi clones. **(B)** Confirmation of TbMyo1 depletion. Whole-cell lysates from three different TbMyo1 RNAi clones were collected at 0, 24, and 48 h and analysed by immunoblotting. The TbMyo1 signal (green) was observed at ~130 kDa, with signal intensity significantly decreasing at 24 h and 48 h in all three clones. Antibodies against TbPFR1,2 were used as loading controls to check for equal sample loading. **(C)** TbMyo1 is essential for *in vitro* growth. The cell population density of TbMyo1 RNAi cells was measured over 96 hours under control (-tet) and induced (+tet) conditions. Cells were diluted after 48 hours to maintain logarithmic growth. Results from two independent experiments, each using three separate clones, are shown; data points represent the mean +/- standard deviation. **(D)** TbMyo1 depletion results in ultrastructural changes in endosomal membranes. BSF TbMyo1 RNAi cells were incubated for 24 h to induce TbMyo1 depletion, followed by high-pressure freezing, embedding in Epon, and examination via transmission electron microscopy. Most cells exhibited either a normal ultrastructure (I) or an extremely enlarged flagellar pocket (V and VI). A subset of cells displayed numerous large and interconnected tubular profiles, suggestive of a swollen endosomal system. Exemplary cells from a single experiment are shown. Abbreviations: clathrin-coated pit: CCP, clathrin-coated vesicle: CCV, endosomal membranes: E, flagellar pocket: FP, Golgi apparatus: G, mitochondrion: M, nucleus: N.
